# Proton FLASH Exposure Preserves Gut Commensal Microbiomes and Spares Intestinal Stem Cells

**DOI:** 10.1101/2025.09.28.679072

**Authors:** Rishi Man Chugh, Jufri Setianegara, Emily Schueddig, Yuting Lin, Payel Bhanja, Shujah Rehman, Stacey Krepel, Devin Koestler, Ronald C Chen, Katherine L Cook, Subhrajit Saha

## Abstract

Emerging evidence shows Proton FLASH radiotherapy can spare normal tissues while maintaining anti-tumor efficacy. However, its impact on intestinal stem cell (ISC) populations and the gut microbiome remains unclear. This is critical, as the gut microbiome influences ISC radiosensitivity. In a mouse model of radiation-induced gastrointestinal syndrome, FLASH-irradiated mice exhibited better survival and less crypt-villus damage compared to mice exposed to conventional proton irradiation. Using scRNA-sequencing, we demonstrated that proton FLASH exposure using pulsed pencil beam scanning spares two distinct ISC populations—*Lgr5+* CBCs and a *Clu+, Mif+, Fabp2+, Anxa2+* revival stem cell (revSC) population—by modulating oxidative stress and cell cycle progression. Analysis of alpha and beta diversity demonstrated that FLASH modulates gut microbiota composition without compromising overall species richness. Notably, FLASH-irradiated mice had higher abundances of *Alistipes* sp. and *Akkermensia* sp., both known for protective effects on ISCs and the intestinal mucosa. The critical role of microbiome in FLASH-mediated sparing effect against radiation toxicity was further confirmed by fecal microbiota transplantation, where FLASH-donor microbiota demonstrated reduced lethality in recipients exposed to proton irradiation with conventional dose rate. Our findings highlight the crucial role of the microbiome in the FLASH-mediated sparing of the mucosal epithelium.

## 1. Introduction

Acute gastrointestinal (GI) toxicity significantly limits the effectiveness of abdominal radiotherapy ^[1]^. Patients with abdominal malignancies often experience severe (grade 3) mucosal toxicity, potentially leading to insufficient tumoricidal radiation doses, early treatment withdrawal, tumor progression or recurrence, and ultimately, a poorer prognosis [2]. The rapid self-renewal rate of intestinal epithelium makes it highly susceptible to radiation-induced DNA damage, resulting in significant radiosensitivity compared to more slowly self-renewing tissues [3]. Our previous findings suggest that intestinal stem cells, the foundation of epithelial architecture, are key determinants of intestinal epithelium survival following radiation exposure [3, 4].

Dose rate significantly impacts tissue radiosensitivity [5]. Ultra-high dose rate (FLASH) radiotherapy (RT) (≥40-60 Gy/s) can potentially spare normal tissues while maintaining the anti-tumor effect, though this effect can be tissue-dependent [6, 7]. However, the precise biological mechanisms underlying the FLASH effect remain largely unclear. Some studies propose that differential oxygen depletion at ultra-high dose rates contributes to the selective sparing of healthy tissues in FLASH RT [8–10]. Specifically, the rapid depletion of oxygen during FLASH RT may minimize the toxic biological effects of free radicals compared to conventional X-ray radiation by increasing the likelihood of radical-radical recombination due to a transient, high concentration of radicals during a short timeframe [9]. Here, we investigated the biological effects of proton FLASH delivered with the pulsed pencil beam scanning IBA Proteus®ONE system, reported for the first time in this study. ISC hierarchy plays a significant role in intestinal epithelial homeostasis and repair [11, 12]. Previous reports suggested that FLASH RT preserves different ISC lineages [7, 13].

However, the mechanism of FLASH effect on ISCs is not clear. Moreover, FLASH effect on ISC regenerative response, cell cycle and oxidative stress has not been studied.

Intestinal microbiomes play a significant role in gut health [14]. Radiation-induced oxidative stress and ROS production in intestinal epithelium can create an unfavorable environment for the survival of anaerobic commensal bacteria, exerting a selection pressure that reduce their populations and promote dysbiosis, thereby disrupting the delicate balance of the gut microbiome [15]. Mucosal dysbiosis substantially contributes to mucosal injury by impairing the regenerative function of intestinal stem cells (ISCs), inducing inflammation, and disrupting the mucosal barrier. Therefore, minimizing mucosal dysbiosis and maintaining a healthy commensal microbiome are crucial for mitigating radiation-induced intestinal injury [16].

However, the effect of radiation dose rate on mucosal dysbiosis has not been well-examined. Given that commensal microbiomes are primarily anaerobic, the radiation-induced production of oxidative radicals may influence the commensal microbiome. Considering the role of FLASH radiotherapy in regulating free radical production, the effect of Proton FLASH on the gut microbiome needs to be investigated in a murine model of abdominal irradiation.

This manuscript demonstrates that delivering a lethal dose of radiation in FLASH mode preserves the gut commensal microbiome with the enrichment of mucoprotective *Alistipes* sp., spares intestinal stem cell lineages, and reduces overall lethality compared to conventional dose rates.

## 2. Result

### 2.1 Proton FLASH reduces severe morbidity compared with Proton conventional dose rate

To evaluate the therapeutic advantage of Proton FLASH over conventional (CONV) proton irradiation in sparing normal intestinal epithelium, a survival study was performed in a mouse model of radiation-induced intestinal injury [3, 4]. Mice were subjected to abdominal irradiation at three different doses: 14 Gy, 16 Gy, and 18 Gy. For Proton FLASH, two dose rates were used: low dose rate (LDR) (43.21–49.38 Gy/sec) and high dose rate (HDR) (57.14–64.29 Gy/sec). This was compared to CONV proton irradiation delivered at a standard dose rate of 0.5–1.0 Gy/sec (Figure 1 A, B). Survival and body weight were monitored over 40 days to analyze the acute toxicity and post-irradiation recovery.

**Figure 1:**
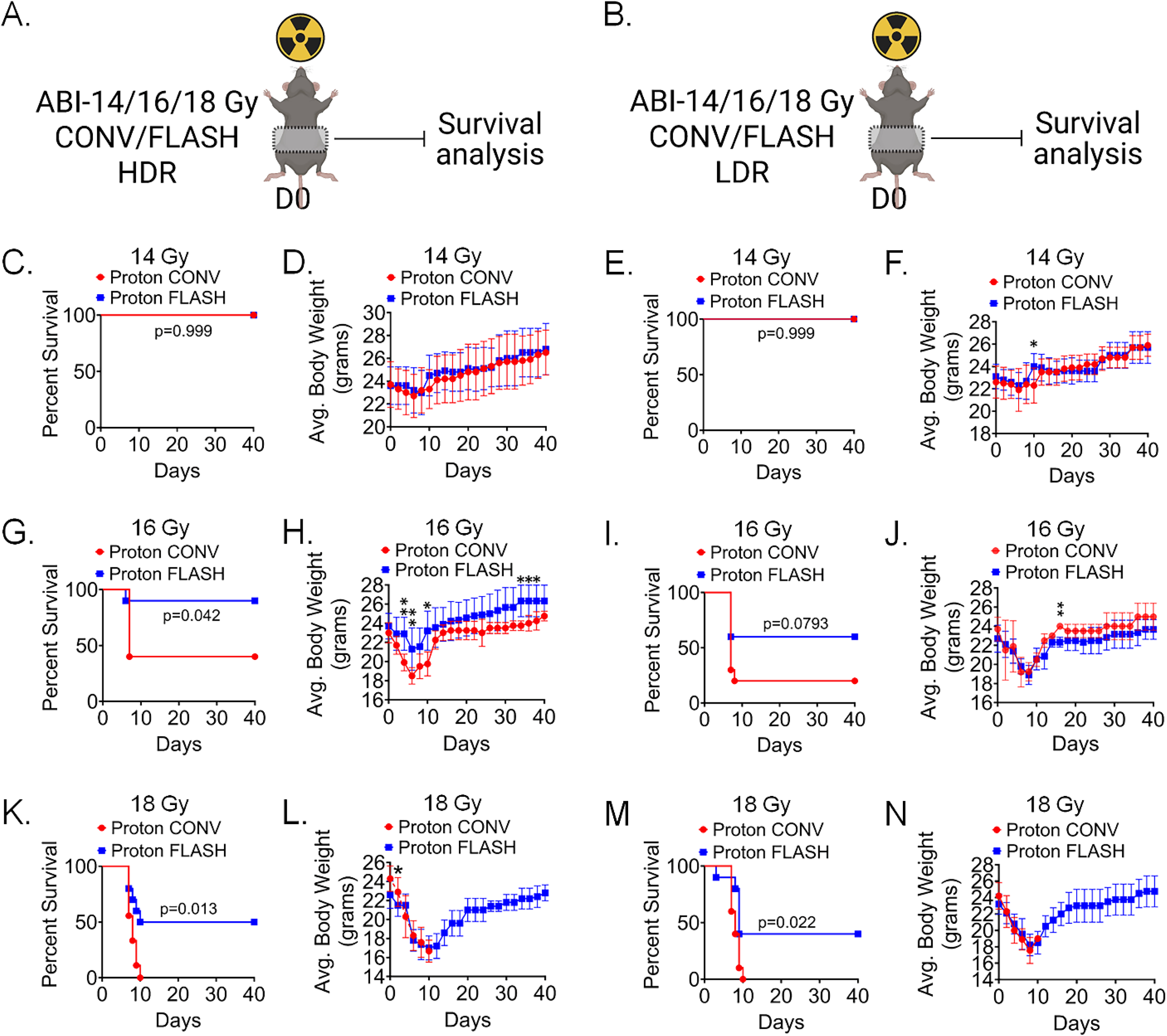
Proton FLASH irradiation reduces morbidity compared with Proton conventional dose rate. A-B. Schematic representation of the experimental design. Mice were exposed to ABI at doses of 14, 16, or 18 Gy using either Proton FLASH or CONV proton irradiation at two dose-rate ranges: low-dose rate (LDR; 43.21–49.38 Gy/s) and high-dose rate (HDR; 57.14–64.29 Gy/s), while CONV proton irradiation was delivered at a standard clinical dose rate of 0.5–1.0 Gy/s. In response to 14 Gy ABI Kaplan-Meier survival curves

At 14 Gy, both FLASH and CONV proton irradiation demonstrated 100% survival under both HDR and LDR conditions, indicating minimal toxicity induced by the 14 Gy dose regardless of dose rate (Figure 1 C, E). Consistent with the survival data, there were no significant differences in body weight between the groups (Figure 1 D, F).

At a radiation dose of 16 Gy, the survival advantage of Proton FLASH over CONV proton irradiation became distinctly evident. Under HDR conditions, mice irradiated with Proton FLASH exhibited a significantly improved survival rate of 90%, compared to only 40% in the CONV proton group (Figure 1 G). A similar trend was observed under low-dose rate (LDR) conditions, where Proton FLASH showed 60% survival, compared to 20% survival with CONV proton irradiation (Figure 1 I). The body weight analysis further supported these findings, as mice irradiated with Proton FLASH demonstrated more rapid recovery of body weight compared to those exposed to CONV proton irradiation under HDR conditions. The average body weight gain was significantly higher in mice irradiated with Proton FLASH after 30 days of irradiation, indicating enhanced post-irradiation recovery (Figure 1 H).

*(C & E) and longitudinal body weight (D & F) of mice demonstrated no differences in FLASH (HDR and LDR) vs CONV. In response to 16 Gy ABI both FLASH-HDR & LDR demonstrated improved survival (90% & 60%) (p=0.042, p=0.0793) (G & I). and body weight (H, J) compared to CONV-irradiated mice. Exposure to 18 Gy ABI, CONV irradiation which resulted in 100% mortality compared to improved survival with FLASH (HDR & LDR) (50% & 40% respectively) (p=0.013, p=0.022.) (K, M). Body weight gain was not comparable due to 100% lethality in conventional proton exposure with 10 days (L, N). Data are presented as Mean ± SD; with Significant; *p < 0.05, **p < 0.005, n=10 mice per group*.

At the highest irradiation dose of 18 Gy, the survival benefit of Proton FLASH irradiation was even more pronounced. Conventional (CONV) proton irradiation resulted in 100% mortality under both HDR and LDR conditions (Figure 1 K, M). In contrast, Proton FLASH irradiation demonstrated 50% and 40% survival rates under HDR and LDR conditions, respectively (Figure 1 K, M). Corresponding body weight data for mice irradiated at 18 Gy showed a gradual recovery following initial loss with proton FLASH, while proton CONV irradiation led to a decline (Figure 1 L, N). These findings suggest that proton FLASH irradiation has the potential to mitigate acute radiation-induced gastrointestinal toxicity and enhance survival after abdominal irradiation.

### 2.2 Proton FLASH irradiation effectively preserves intestinal morphology and enhances crypt-villus structure compared to CONV

Given the tissue-specific sensitivity of the gastrointestinal (GI) tract to radiation-induced injury, this study next investigated intestinal epithelial morphology in response to Proton FLASH CONV Proton irradiation.

To assess the impact of high-dose (18-21 Gy) abdominal irradiation (ABI) on intestinal epithelium, histological analyses of jejunal sections were conducted 72 hours post-irradiation with either high dose rate (HDR) or low dose rate (LDR) proton FLASH or CONV Proton irradiation (Figure 2 A, Supplementary Figure 2 A). Histological evaluation revealed that proton FLASH therapy significantly better-preserved intestinal morphology than CONV irradiation, under both 18 Gy and 21 Gy doses with HDR and at 18 Gy under LDR condition (Figure 2 B, Supplementary Figure 2 B).

**Figure 2:**
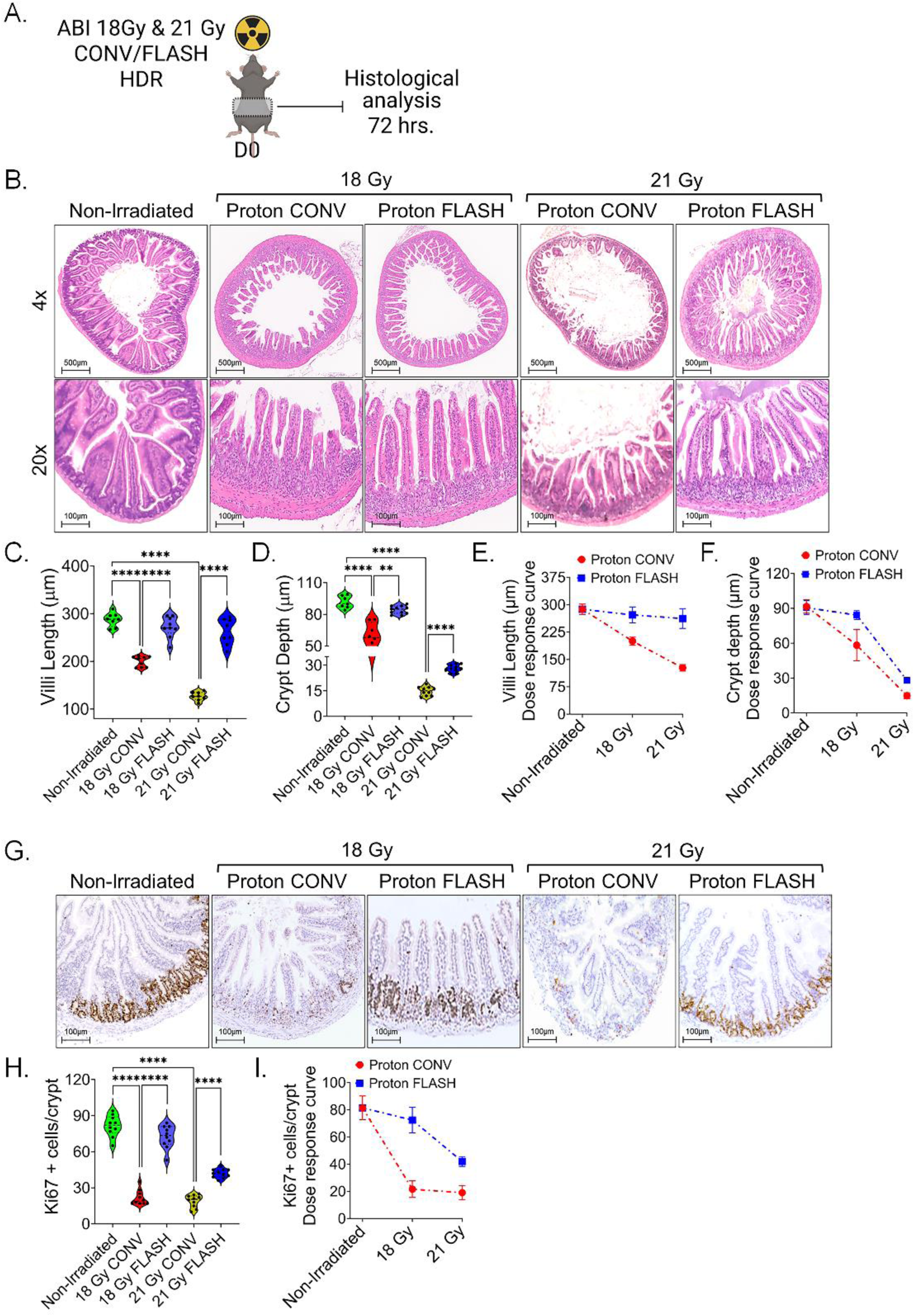
Proton FLASH irradiation preserves intestinal morphology following high-dose ABI. (A) Schematic representation of experimental design. Mice received 18 Gy and 21 Gy abdominal irradiation using either CONV or FLASH proton irradiation under HDR dose rate settings, followed by histological analysis of jejunal sections at 72 hours post-irradiation. (B) Representative H&E-stained jejunal sections (Transverse sections: top panel and longitudinal sections: Bottom panel) from CONV- and FLASH-irradiated mice under HDR conditions show improved mucosal preservation in FLASH-irradiated groups at both 18 Gy and 21 Gy of irradiation. (C, D) Violin plot demonstrates higher villus length and crypt depth in FLASH-irradiated mice at both 18 Gy and 21 Gy of irradiation compared to CONV dose rate. (E, F) Dose-response curves demonstrated less decline in villus length and crypt depth with increasing dose from 18 Gy to 21 Gy in FLASH compared to CONV dose rate. (G, H) showing the number of Ki67+ proliferative cells per crypt. FLASH-irradiated mice demonstrate significantly higher Ki67+ cell counts compared to CONV-irradiated animals at both 18 Gy and 21 Gy of irradiation. (I) The dose-response curve for Ki67+ cells demonstrated FLASH irradiation minimizes the radiation-induced decrease in crypt proliferation. Data presented as mean ± SD, with Significant; **p < 0.005, ****p < 0.00005. n=10 mice per group.

Tissues from FLASH-exposed mice exhibited well-defined villi and intact crypt structures, demonstrating maintained mucosal integrity. In contrast, CONV-irradiated intestinal tissue showed severe radiation-induced damage, including blunted villi, crypt depletion, and mucosal flattening, particularly at higher 21 Gy of radiation dose (Figure 2 B, Supplementary Figure 2 B). Quantitative morphometric analysis further supported these observations, at both 18 Gy and 21 Gy under HDR, FLASH-irradiated mice showed significantly greater villus length and crypt depth compared to CONV-irradiated mice (Figure 2 C, D). Dose-response curves showed a clear decline in villus length and crypt depth with increasing dose from 18 Gy to 21 Gy, but FLASH irradiation consistently reduced this decline relative to CONV irradiation (Figure 2 E, F).

Under LDR conditions with 18 Gy, FLASH irradiation also significantly improved both villus length and crypt depth relative to CONV irradiation, demonstrating that its protective effects extend across different dose rates (Supplementary Figure 2C–D).

To further assess epithelial regenerative potential, Ki67 immunohistochemistry was used to quantify proliferating crypt cells. At 18 Gy, FLASH-irradiated mice (under both HDR and LDR) showed a markedly higher number of Ki67+ cells per crypt compared to CONV-irradiated mice (p < 0.00005) (Figure 2 F, G and Supplementary Figure 2 B, E). Under HDR condition, Ki67+ cell numbers declined with doses in both groups, but FLASH-irradiated mice retained significantly greater proliferative capacity than CONV-irradiated mice at both 18 Gy and 21 Gy.

The dose-response curve for Ki67+ cells under HDR confirmed this pattern, FLASH irradiation preserves the radiation-induced decline in crypt proliferation more effectively than CONV irradiation (Figure 2 H). Collectively, these results provide strong histopathological evidence that proton FLASH irradiation is more effective than CONV proton irradiation in sparing intestinal epithelium from radiation toxicity.

### 2.3 Proton FLASH preserves the Intestinal Stem Cell population

Regenerative response of intestinal epithelium primarily depends on Lgr5+ highly self-renewing intestinal stem cells (ISCs) also known as crypt base columnar cells (CBCs) [3, 4, 17, 18]. To investigate whether Proton FLASH preserves this ISC population, Lgr5-EGFP-CreERT2; R26-ACTB-tdTomato-EGFP reporter mice were used [18]. These mice enable direct visualization of Lgr5+ ISCs within the intestinal crypts. Mice were irradiated with Proton FLASH and CONV 21 Gy ABI and intestinal tissues were isolated 96 hours post-irradiation for analysis (Figure 3 A). Analyzing Lgr5+ ISCs at 96 hours, following the apoptotic phase at 48-72 hours, provides accurate surviving fractions for intestinal epithelial regeneration. Confocal imaging of jejunal sections showed a marked depletion of Lgr5+ ISCs in CONV-irradiated mice, characterized by a significant reduction in GFP+ (Green fluorescent Protein) crypts per field (Figure 3 B, C) when compared to non-irradiated and FLASH-irradiated mice (p<0.0005 and p<0.005, respectively). No significant difference was found in the density of Lgr5+ cells in FLASH-irradiated mice compared to non-irradiated mice, suggesting preservation of the functional regenerative niche (Figure 3 B, C). These findings demonstrate that in addition to maintaining crypt-villus morphology, Proton FLASH irradiation also preserves the intestinal stem cell pool following high-dose irradiation, thereby supporting tissue homeostasis and post-injury regeneration.

**Figure 3:**
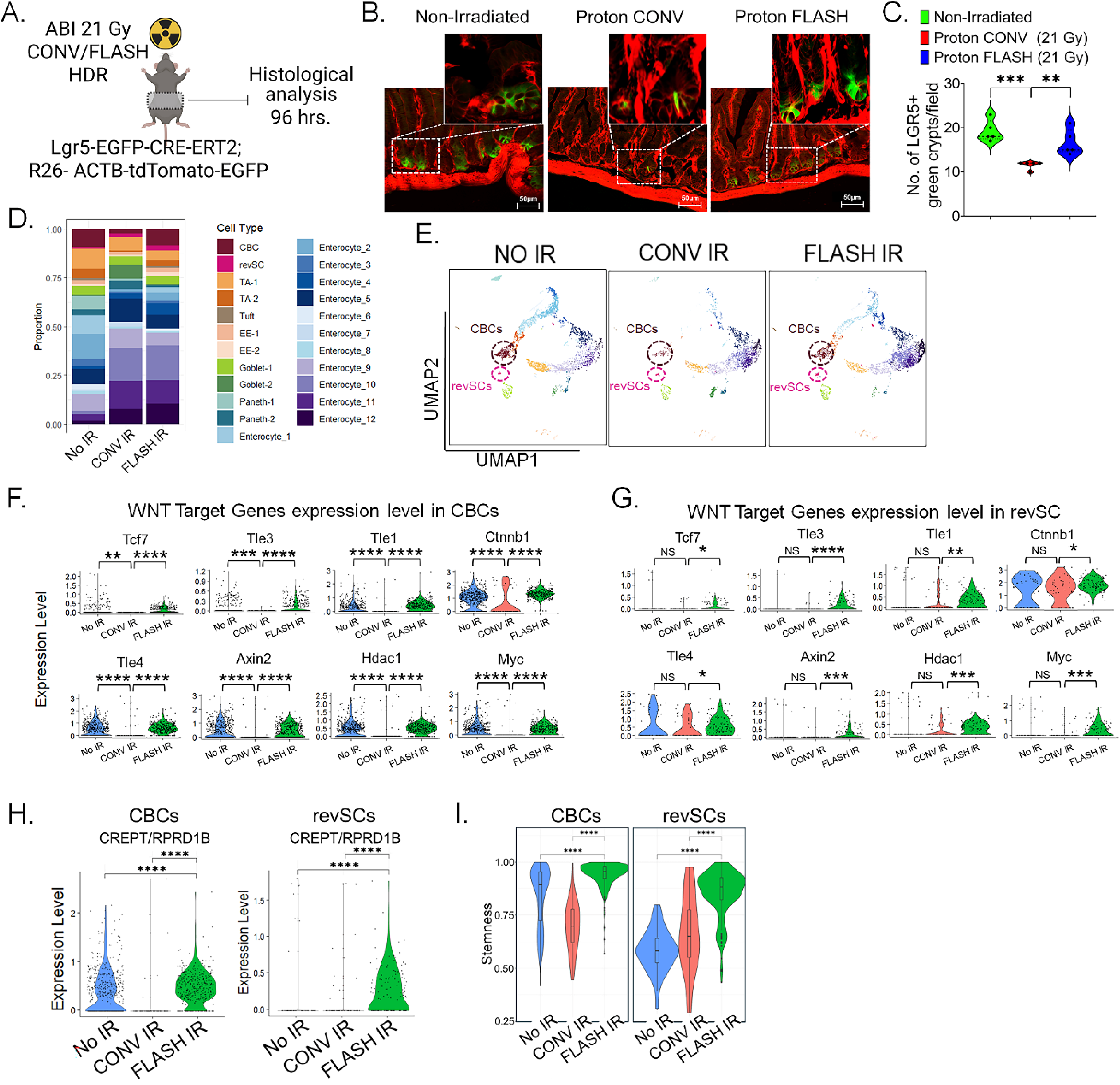
Proton FLASH spares intestinal stem cell population following high-dose abdominal irradiation. (A) Schematic representation of experimental design using Lgr5-EGFP-CreERT2; R26-ACTB-tdTomato-EGFP reporter mice exposed to 21 Gy ABI using HDR FLASH or CONV proton delivery, followed by tissue collection at 96 hours post-irradiation for histological analysis. (B) Representative confocal images of jejunal sections from non-irradiated, CONV, and FLASH-irradiated mice showing Lgr5+ intestinal stem cells (green) and tdTomato+ (red). Insets highlight higher magnification views of Lgr5+ crypts. (C) Violin plot demonstrates quantification of Lgr5+ crypts per field shows significantly higher stem cell preservation in FLASH-irradiated mice compared to CONV-irradiation. (D) Bar plot showing the proportional representation of each intestinal epithelial cell type analyzed by scRNA seq in No IR, CONV IR, and FLASH IR groups. Higher presence of crypt base columnar cells (CBCs) and revival stem cell (revSCs) was noted in FLASH IR group compared to CONV IR. (E) UMAPs representing epithelial cell cluster distribution in: No IR, CONV IR and FLASH IR. Enrichment in cluster representing CBCs and revSCs population was noted. (F-G) violin plot demonstrated higher Wnt target genes expression levels in CBCs (F) and in revSCs (G) in FLASH IR compared to CONV IR group. (H, I) Violin plot demonstrated higher CREPT/RPRD1B expression level and stemness score in FLASH irradiated CBCs and revSCs group compared to CONV IR group. (Significant; *p<0.05, **p<0.005, ***p < 0.0005, ****p < 0.00005, NS: not significant), n = 3 mice per group.

### 2.4 Proton FLASH spares ISC lineages and other intestinal epithelial cell population

To investigate the regenerative and differentiation dynamics of the intestinal epithelium after irradiation, single-cell RNA sequencing (scRNA-seq) was used to characterize epithelial cell populations. This was performed on small intestinal cells isolated 72 hours after 21 Gy of ABI using CONV-irradiated, FLASH-irradiated and non-irradiated mice. Single-cell analysis of intestinal epithelium revealed 24 distinct epithelial subpopulations, indicative of specialized lineages and varying maturation stages. Initial clustering of integrated samples established a comprehensive dataset of mouse intestinal cells. Epithelial cells, identified by Epcam expression, were then isolated and re-integrated for further analysis (Supplementary Figure 3 A, B). Subsequent clustering revealed 24 clusters with unique gene expression profiles that defined functional identities (Supplementary Figure 3 C) of different epithelial cell types. Two distinct intestinal stem cell populations were identified: actively cycling crypt base columnar (CBC) stem cells (marked by Smoc2, Lgr5, and Ascl2 expression) [17, 19] and injury-induced revival stem cells (revSCs) (characterized by expression of Ly6a, Clu, Areg, and Anxa2), previously linked to tissue regeneration following injury [20, 21]. Proportional bar plots and UMAPs showing the absolute number of cells demonstrated that CONV irradiation depleted CBCs and revSCs (p=0.2546) compared to FLASH irradiation. (Figure 3 D, E). In contrast, FLASH irradiation preserved CBCs at levels comparable to non-irradiated controls (p=0.9108) and significantly increased revSCs (p=0.0017). Collectively, these findings indicate that Proton FLASH (not only) preserves crypt-villus architecture and Lgr5⁺ ISCs but also maintains a diverse and functional pool of regenerating epithelial progenitors crucial for mucosal recovery post-irradiation.

### 2.5 Enhanced Wnt Signaling in ISCs populations following proton FLASH irradiation

Wnt signaling plays an important role in maintaining ISCs homeostasis, proliferation, and regeneration for epithelial renewal following radiation-induced injury [17, 22]. To investigate whether FLASH irradiation preserves or enhances Wnt signaling in ISCs compared to CONV irradiation, we examined the expression of Wnt target genes in CBCs and revSCs. In both CBCs and revSCs FLASH-irradiated cells demonstrated significantly higher expression of canonical Wnt target genes including *Axin2*, *Myc*, *Tcf7*, *Ctnnb1*, and *Tcf7l2*, compared to CONV-irradiation (Figure 3 F, G). Additionally, in CBCs and revSCs the upstream transcriptional regulators such as *Tcf7*, *Tcf7l2* and corepressors like *Tle1*, *Tle3*, and *Tle4* were also highly expressed in FLASH-irradiated cells compared to CONV-irradiated cells, further supporting the activation of the Wnt/β-catenin signaling.

These findings suggest that FLASH irradiation not only spares but actively supports the regenerative capacity of ISCs by enhancing Wnt signaling compared to CONV irradiation. To further explore the downstream effects of enhanced Wnt/β-catenin signaling in FLASH-irradiated ISCs, we assessed the expression of CREPT/RPRD1B (Cell-cycle Related and Expression-elevated Protein in Tumor), a transcriptional co-regulator known to promote cell proliferation by enhancing Wnt/β-catenin signaling [23]. CREPT/RPRD1B has been involved in supporting epithelial regeneration by enhancing ISCs regenerative capacity and maintaining stem cell identity following radiation injury [24, 25]. Radiation-induced injuries often impair stemness of intestinal epithelial cells. To evaluate the impact of proton FLASH irradiation on ISCs, we analyzed CREPT/RPRD1B expression and stemness scores measured by CytoTRACE algorithm [26] in CBCs and revSCs (Supplementary Figure 4 A, B). CREPT/RPRD1B expression was significantly higher in both CBCs and revSCs of FLASH-irradiated cells compared to CONV-irradiated cells (Figure 3 H). Remarkably, both CBCs and revSCs also demonstrated significantly higher stemness scores in FLASH-irradiated cells, which suggest a preservation or enhancement of their stem cell identity (Figure 3 I). In contrast, CONV irradiation led to a significant reduction in stemness in both CBCs and revSCs (Figure 3 I). These results further indicate that proton FLASH irradiation promotes CREPT/RPRD1B expression and sustains stemness in ISCs populations which potentially contributing to epithelial regeneration and reduced radiation-induced toxicity.

Further analysis of regenerative response of revSCs to proton FLASH irradiation multiple pathways were significantly upregulated, associated with stem cell survival, epithelial repair, and tissue regeneration (Figure 4 A, B). Among the top enriched pathways were TGF-β receptor signaling and SMAD-mediated transcriptional activation, which are known to play critical roles in epithelial homeostasis, regeneration, and modulation of the stem cell niche [27, 28]. Activation of TGF-β signaling is essential for balancing ISC proliferation and differentiation and has been implicated in protecting the intestinal barrier during injury repair [28]. We also observed an enrichment in Hedgehog signaling-related pathways such as hedgehog ligand biogenesis and signaling by Hedgehog. Hedgehog signaling is known to support intestinal regeneration by influencing mesenchymal niche activity and maintaining epithelial architecture [29].

**Figure 4:**
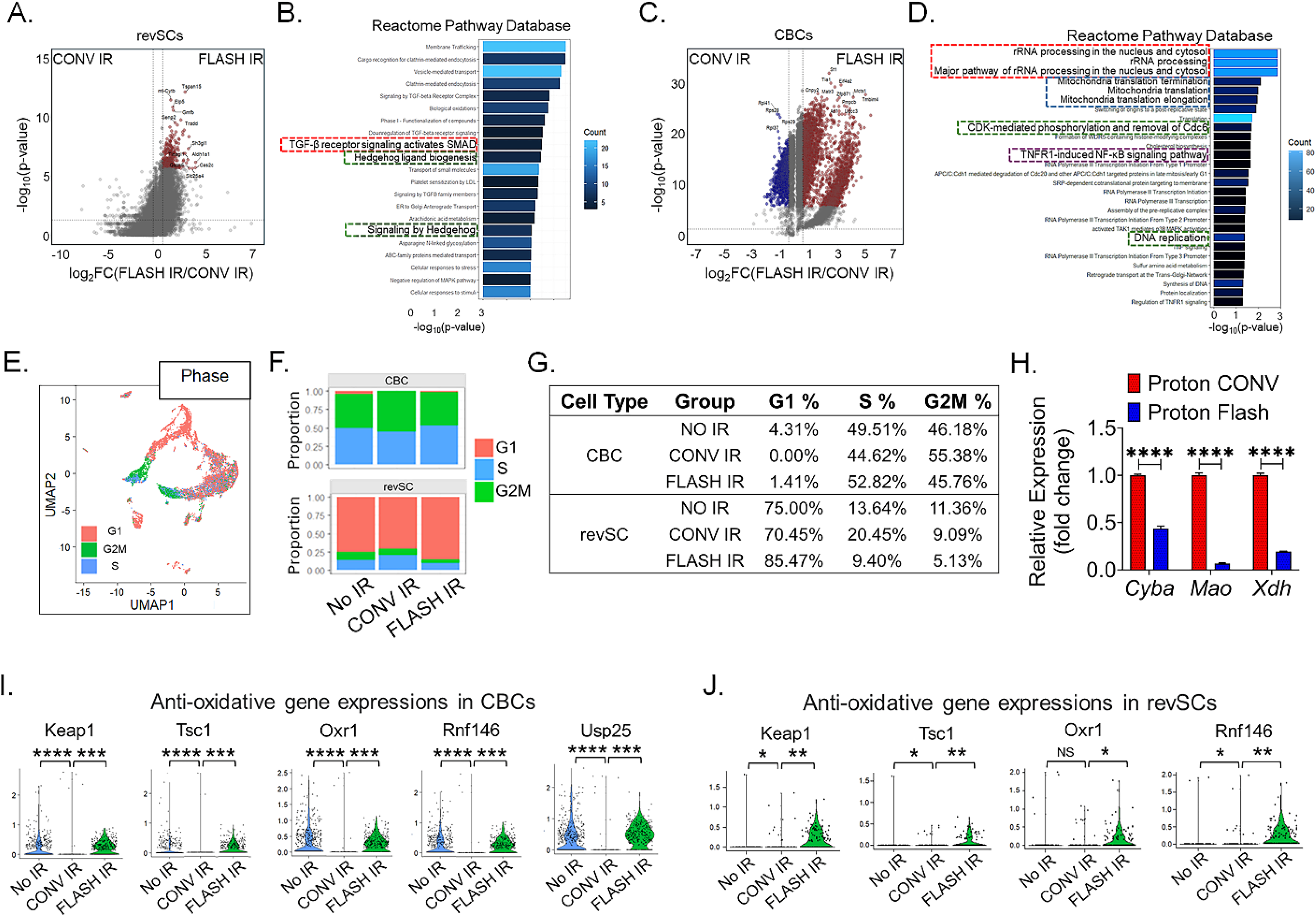
Proton FLASH irradiation activates regenerative pathways in CBCs and revSCs. (A, B) Pathways enriched in FLASH-irradiated revSCs include TGF-β receptor/SMAD signaling, Hedgehog signaling, and other epithelial repair pathways, indicating enhanced stem cell survival, niche modulation, and barrier protection. (C, D) Pathway analysis of genes upregulated in CBCs following FLASH irradiation showing significant enrichment in pathways supporting the rapid proliferation and regenerative function of ISCs after radiation injury (red box, blue box), restoring mucosal integrity (green box) and modulating inflammatory responses and promoting survival signaling (purple box). (E) UMAP showing cell cycle phase distribution (G1, S, G2/M) across epithelial cells. (F) Bar diagram demonstrating quantification of cell cycle phases (G1, S, G2/M) within CBC, revSC clusters across different groups. (G) Table demonstrating cell cycle phase distribution of selected intestinal epithelial cell types (CBC, revSC). (H) qPCR analysis demonstrated, low expression of oxidative stress–related gene expression in FLASH irradiated intestinal epithelial cells compared to CONV irradiated cells. (I, J) scRNA-seq analysis showing significant upregulation of antioxidant genes (Keap1, Usp25, Tsc1, Oxr1, Rnf146) in CBCs and revSCs following FLASH irradiation, indicating enhanced oxidative stress defense. (Significant; *p<0.05, **p<0.005, ***p < 0.0005, ****p < 0.00005, NS: not significant).

To identify molecular mechanisms underlying the regenerative advantage of proton FLASH irradiation, a pathway analysis of genes upregulated in CBCs was performed. Several pathways were significantly upregulated in CBCs following FLASH irradiation (Figure 4 C, D), among the most significantly enriched pathways were rRNA processing and mitochondrial translation, which are essential for ribosome biogenesis and protein synthesis. These pathways play a significant role in supporting the rapid proliferation and regenerative function of ISCs after radiation injury [30]. The Pathways involved in cell cycle regulation and replication licensing, such as CDK-mediated phosphorylation and removal of Cdc6 and DNA replication, were also significantly upregulated. These pathways are required for proper progression through the G1/S phase and initiation of DNA replication [31], suggesting that FLASH irradiation enables ISCs populations to more efficient re-entry into the cell cycle following radiation-induced damage for restoring mucosal integrity. TNFR1-induced NF-κB signaling pathways are known to play protective and regenerative roles in the intestinal epithelium [32]. NF-κB activation in ISCs and niche cells promotes epithelial regeneration by modulating inflammatory responses and promoting survival signaling [33].

### 2.6 FLASH regulates expression of cell cycle related genes in ISCs

Analysis of cell cycles related genes in CBCs and revSC suggested that FLASH exposure delayed the cell cycle progression post exposure compared to conventional exposure. At 72 hours post CONV exposure, a higher percentage of CBCs progressed to G2/M phase (55.38%) compared to FLASH (45.76%) (Figure 4 E-G). Similar response was also observed in revSCs. Due to quiescence and slow cycling nature, a majority of the revSCs were observed at G1 phase and very minimum number of cells progressed to G2/M phase (Figure 4 E-G). However, revSCs with CONV exposure had a higher percentage of cells in S and G2/M phase. (Figure 4 G). It has been reported that transient cell cycle arrest and thereby delay in cell cycle progression immediately following radiation in ISCs helps to reduce DNA damage [34]. Our data further suggested that delay in progression of cell cycle following FLASH exposure may contribute to preservation of CBCs and revSCs compared to CONV exposure.

Collectively, these findings suggest that proton FLASH irradiation triggers a distinct transcriptional program in CBCs that enhances cell survival, facilitates re-entry into the cell cycle and promotes regenerative signaling and activation of revSCs which enables more efficient regeneration of the intestinal epithelium compared to CONV irradiation.

### 2.7 Proton FLASH exposure modulates oxidative stress in ISCs

It has been reported that normal-tissue sparing effect of FLASH-RT is due to the rapid reduction of intracellular oxygen during FLASH-RT, namely radiolytic oxygen depletion (ROD) [8, 35]. Oxygen increases the radiation susceptibility of cells with a maximum OER between 2.5 and 3.5 [36]. Compared to CONV RT, FLASH RT dose rate promote depletion of O2 critical to the biological response due to competition between the depletion rate and the re-diffusion rate of O2 from the capillaries [36]. FLASH may increase the probability of radical–radical recombination and therefore minimize the toxic biological effect of free radicals such as oxidative stress due to transient accumulation of a large concentration of radical species during a short time window [37]. In mice model of abdominal irradiation, we have observed reduced expression of oxidative stress genes such as *Cyba, Mao, Xdh* (Figure 4 H) in intestinal epithelium with exposure to Proton FLASH compared to CONV at 72 hours post exposure. scRNA seq analysis demonstrated significant upregulation of multiple anti-oxidative gene expressions such as *Keap1, Usp25, Tsc1, Oxr1, Rnf146* in CBCs (Figure 4 I) and revSCs (Figure 4 J) in response to FLASH irradiation compared to CONV irradiation.

These findings point to a dual mechanism for FLASH irradiation’s tissue-sparing effects in mice intestinal tissue: controlling tissue injury stress responses while simultaneously maintaining regenerative signaling.

### 2.8 Proton FLASH Irradiation preserves gut microbiome diversity and composition

Proton FLASH irradiation preserves gut microbial diversity more effectively than CONV irradiation, although differences in community composition remain significant compared to non-irradiated controls. To evaluate the impact of irradiation dose on the intestinal microbiome, we analyzed fecal samples collected from mice irradiated with either Proton FLASH or CONV irradiation. Alpha diversity analysis, using Chao1 index, revealed that all CONV irradiated groups demonstrated increased richness relative to non-irradiated controls. FLASH-irradiated mice maintained significantly higher microbial diversity compared to the non-irradiated controls, only at 72 hours post-irradiation and when compared with CONV irradiation at 72 hours was significantly decreased (Figure 5A). Shannon α-diversity index measurement indicates CONV irradiation, but not FLASH, resulted in elevated α-diversity (Figure 5B). Beta diversity analysis using Bray-Curtis Principal Coordinate Analysis (PCoA) demonstrated a distinct clustering of microbial community between FLASH and CONV irradiated groups. Further the PERMANOVA analysis confirmed statistically significant differences between these distinct clustering (Figure 5 C), which suggests that the Proton FLASH and CONV, two irradiation modalities, induce markedly different effects on gut microbial composition. Phyla taxonomic profiling showed that both radiation modality reduced Bacillota phyla proportional abundance; However, CONV irradiation resulted in a higher proportional abundance when compared with FLASH irradiation at matching timepoints (Figure 5D, E), whereas FLASH IR displayed higher proportional abundance of Verrucomicrobiota and lower Bacillota (Figure 5 D). Both CONV and FLASH irradiation resulted in an increase Verrucomicrobiota abundance when compared to non-irradiated controls. At 24 hours and 72 hours post irradiation, FLASH led to significantly higher proportional abundance of Verrucomicrobiota than CONV (Figure 5 D, E). These findings suggest that both modalities of irradiation shift the gut microbiome; However, CONV irradiation modifies microbial richness and evenness more severely than FLASH irradiation, suggesting FLASH irradiation may better preserve gut microbial community structure and diversity.

**Figure 5:**
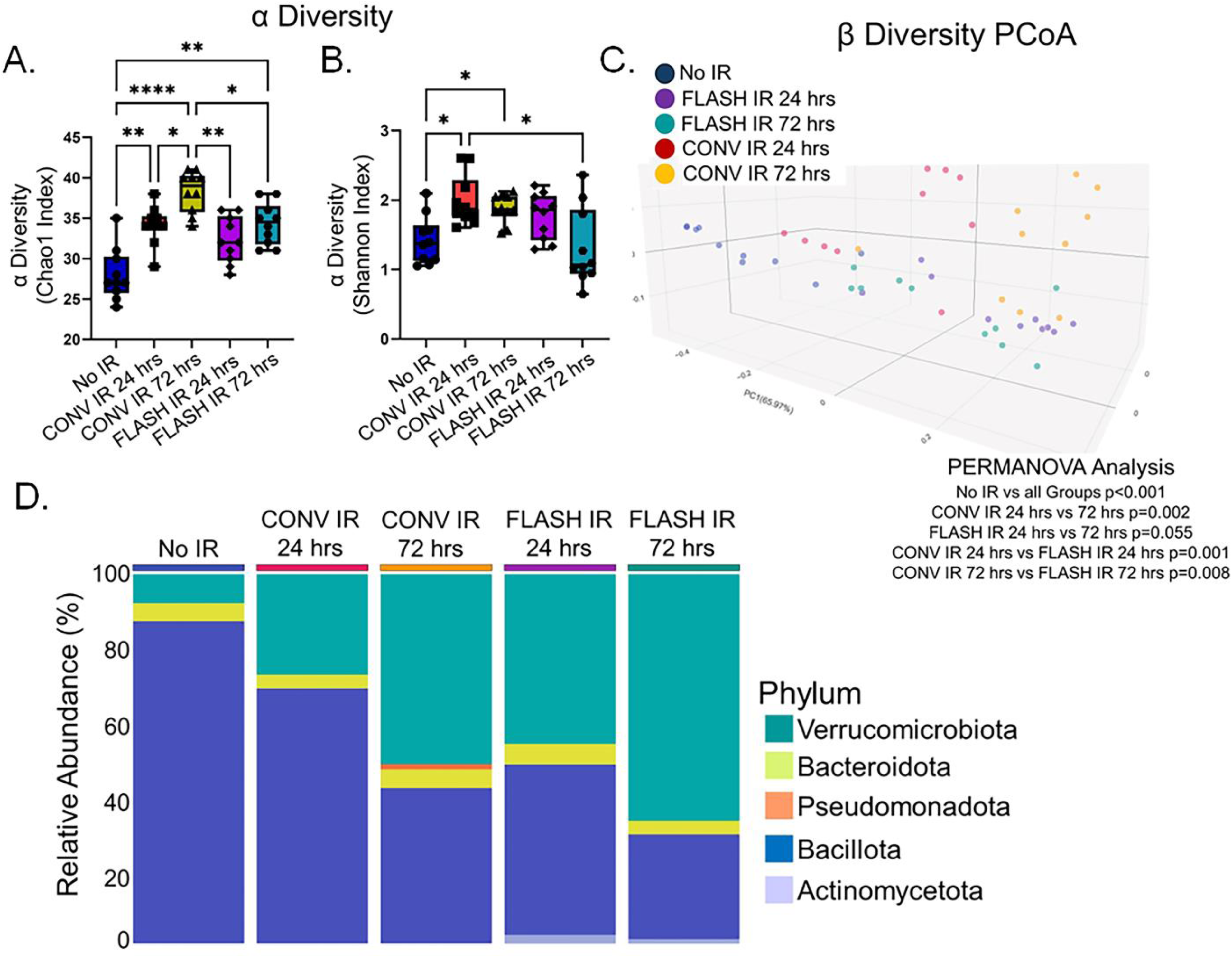
Proton FLASH irradiation preserves gut microbial diversity and composition compared to CONV irradiation. (A) Alpha diversity analysis (Chao1 index) showing that all CONV-irradiated groups exhibited increased richness relative to non-irradiated controls. FLASH-irradiated mice maintained significantly higher diversity only at 72 h post-irradiation compared to controls, but diversity was significantly decreased relative to CONV at the same timepoint. (B) Shannon α-diversity index demonstrates that CONV, but not FLASH, irradiation resulted in elevated α-diversity. (C) Beta diversity analysis using Bray–Curtis Principal Coordinate Analysis (PCoA) showing distinct clustering of microbial communities between FLASH and CONV irradiated groups, with PERMANOVA confirming statistically significant differences. (D) Phylum-level taxonomic profiling indicating that both irradiation modalities reduced Bacillota abundance; however, CONV irradiation caused a marked increase in Bacillota relative to FLASH at matched timepoints, whereas FLASH maintained relatively higher Verrucomicrobiota and lower Bacillota at both 24 and 72 h post-irradiation compared to CONV. . Data are shown as mean ± SD, (Significant; *p<0.05, **p < 0.005, ****p < 0.00005), n = 10 mice per group.

Further, the taxonomic profiling at the species level revealed a distinct microbial composition across groups, with notable enrichment of specific taxa in the FLASH irradiated group (Figure 6A). Among these, *Akkermansia muciniphila* was significantly elevated in FLASH irradiated group at 24 hours compared to CONV irradiated groups suggesting improved mucosal resistance following irradiation [38, 39] (Figure 6B). Similarly, *Schaedlerella arabinosiphila*, *Adlercreutzia equolifaciens*, *Adlercreutzia caecimuris* and *Adlercreutzia mucosicola* were significantly enriched in FLASH irradiation group at various timepoints compared to CONV irradiation groups (Figures 6 C–F), potentially indicating restoration of key short-chain fatty acid and equol-producing commensals which would help in gut barrier integrity and anti-inflammatory properties respectively [40–42]. *Alistipes sp. An31A*, which is associated with gut barrier function and metabolic health, while *Bifidobacterium pseudolongum* which increases the mucin production and strengthens the mucus layer [43] were markedly increased in FLASH irradiated mice compared to CONV irradiation (Figure 6G, H) further supporting the enrichment of beneficial commensal taxa in Proton FLASH irradiation group. In contrast, CONV radiation led to enrichment of *Faecalibaculum rodentium*, *Anaerotruncus sp. G3(2012), Romboutsia timonensis* when compared with non-irradiated controls and FLASH irradiation at various time points (Figures 6 I–K). Previous literature associated GI distress and inflammation with intestinal *Romboitsia timonensis* abundance [44], highlighting microbiota disruption under CONV irradiation. Furthermore, the functional microbial gene enrichment alterations, LEfSe analysis (Figure 6 L) of MetaCyc pathways comparing FLASH irradiation and CONV irradiation at 24 hr revealed significant enrichment of microbial biosynthetic pathways in the FLASH group, including NAD de novo biosynthesis, L-methionine biosynthesis, and folate transformations, suggesting enhanced metabolic potential, redox balance modification [45], and mucosal support [46] in FLASH irradiated mice. Collectively, these data demonstrate that FLASH irradiation preserves beneficial microbial functions.

**Figure 6:**
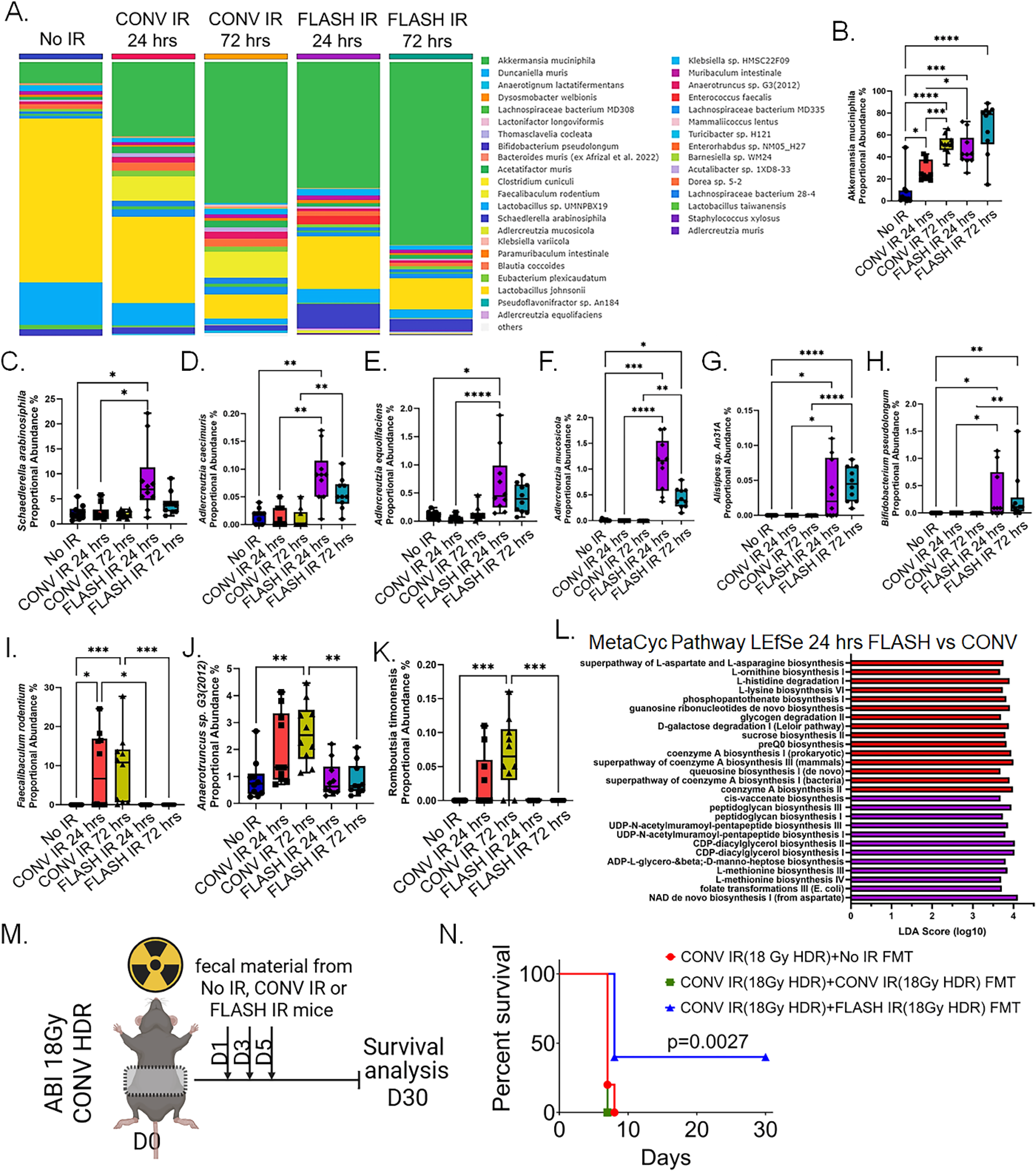
FLASH irradiation preserves beneficial microbial functions. (A) Species-level taxonomic profiling showing distinct microbial composition between groups, with notable enrichment in FLASH-irradiated mice. (B) Significant increase in Akkermansia muciniphila abundance at 24 h post-FLASH irradiation compared to CONV, suggesting improved mucosal resistance. (C–F) Enrichment of Schaedlerella arabinosiphila, Adlercreutzia equolifaciens, Adlercreutzia caecimuris, and Adlercreutzia mucosicola in FLASH mice at both the timepoints, indicating restoration of short-chain fatty acid- and equol-producing commensals that promote gut barrier integrity and anti-inflammatory effects. (G, H) Increased abundance of Alistipes sp. An31A and Bifidobacterium pseudolongum in FLASH mice, taxa associated with enhanced mucin production, mucus layer strengthening, and metabolic health. (I–K) Higher abundance of potentially harmful taxa (Faecalibaculum rodentium, Anaerotruncus sp. G3(2012), Romboutsia timonensis) in CONV irradiation, with R. timonensis previously linked to GI distress and inflammation. (L) LEfSe analysis of MetaCyc pathways at 24 h showing significant enrichment in FLASH mice of biosynthetic pathways including NAD de novo biosynthesis, L-methionine biosynthesis, and folate transformations, suggesting enhanced metabolic potential and mucosal support. (M) Schematic representation of fecal microbiota transplantation (FMT) experimental design in which CONV-irradiated recipients received microbiota from either CONV or FLASH irradiated donor mice. (N) Survival analysis showing 100% mortality within 10 days in CONV recipients of CONV-FMT, versus 40% survival beyond 30 days in CONV recipients of FLASH-FMT, indicating functional protection supported by FLASH-preserved microbiota. (Significant; *p<0.05, **p<0.005, ***p < 0.0005, ****p < 0.00005).

To functionally assess whether the gut microbiota from FLASH exposed mice mitigate radiation toxicity, we conducted a fecal microbiota transplantation (FMT) experiment. The recipient mice were first irradiated to CONV irradiation with LD100/10 (18 Gy, HDR) dose ensuing primarily GI toxicity associated mortality [3, 4, 17, 18] and were then orally gavage with fecal material derived from donor mice that had previously irradiated either CONV or FLASH irradiation (Figure 6 M). CONV irradiated mice receiving FMT from CONV irradiated donors demonstrated 100% mortality with 10-15 days post irradiation suggesting death from radiation induced gastrointestinal syndrome [3, 4, 17, 18]. But FMT from FLASH irradiated mice recued CONV irradiated recipient mice from radiation induced lethality and improved survival in 40% of mice beyond 30 days post irradiation (Figure 6 N). This approach clearly suggested that FLASH mediated enrichment of mucoprotective commensal microbiome such as *Akkermansia muciniphila and Alistipes sp* is beneficial to mitigate intestinal injury. These findings indicate that the microbial communities preserved or enriched by FLASH irradiation are functionally capable of enhancing resistance to radiation-induced injury [47].

## 3. Discussion

Determination of the mechanism behind tissue sparing effect of ultra-high dose rate (FLASH) irradiation is important for its clinical application. In the present study in mice model of radiation induced gastrointestinal syndrome we have observed that proton FLASH irradiation spares intestinal stem cell population including the revSC damage-induced stem cell population with the better preservation of gut microbiome population. The presence of functionally active intestinal stem cells is important to spare mucosal epithelium from radiation injury. FLASH spares intestinal stem cells with the restitution of epithelial crypt villus architectures and minimizes radiation induced lethality in mice.

Exposure to this ultra-high dose rate also demonstrated significant reduction in radiation induced oxidative stress compared to conventional dose rate. All this evidence clearly suggested that FLASH exposure is less damaging to mucosal health compared to conventional proton exposure.

ISC lineages consist of actively self-renewing Lgr5+ve CBCs [20, 48], quiescent Bmi+ve/K19+ve cells located in +4 position [48] from crypt bottom and injury induced revSCs located further higher in crypt [20]. The functional hierarchy of these cell types tightly regulates intestinal epithelial homeostasis and regeneration. In response to genotoxic stress such as radiation loss of these cell types primarily depends on their radio sensitivity. Several reports from our groups clearly demonstrated that Lgr5+ve CBCs are most radiosensitive [3, 4, 49]. Moreover, restitution of these CBCs from radiation toxicity is important to maintain epithelial regenerative response. In mice model of radiation induced gastrointestinal syndrome we observed that FLASH spares the Lgr5+ve CBCs which results maintenance of regenerative epithelium as demonstrated by Ki67+ve cells and TdT+ve cells. Better survival outcomes along with less damage in crypt villus architecture in mice exposed to FLASH compared to conventional dose rate further validate the sparing effect FLASH on ISCs. Revival stem cells (revSCs) defined by transient induction of clusterin (CLU) expression rapidly expand and differentiate into multiple intestinal epithelial lineages during intestinal regeneration [25]. Therefore, presence of functional revSCs is also important to repopulate different cell types of intestinal epithelium post injury. FLASH exposure spares revSCs compared to conventional dose rates suggesting that FLASH is a better option to maintain post irradiation mucosal integrity.

FLASH exposure minimizes normal tissue oxygen consumption which is similar to hypoxic state [50]. We have observed that compared to conventional dose rate, FLASH exposure results less oxidative stress and ROS production suggesting less oxygen consumption in intestinal epithelial cells. scRNA seq data on cell cycle markers also suggested that ISCs including Lgr5+ve CBCs and revSCs are slower in cell cycle progression with FLASH exposure compared to conventional dose rate. At 72 hours post CONV exposure a higher percentage of CBCs and revSCs progressed to G2/M phase compared FLASH. Reduction in cell cycle progression may contribute to less oxygen consumption due to reduction in energy demand [51]. In case of lower energy demand cellular metabolism shifts from aerobic to anerobic/glycolytic state. In the present study scRNA seq data demonstrated significant upregulation of glycolysis associated gene expression with FLASH exposure compared to CONV exposure. Slower cell cycle progression also makes these ISCs less vulnerable to radiation induced DNA damage compared to conventional exposure. In summary FLASH exposure supports anerobic metabolism in ISCs resulting less oxidative stress. Lower metabolic demand due to slower cell cycle progression in ISCs following FLASH exposure further contribute to this metabolic shift.

Commensal gut microbiota is primarily anaerobic and very much sensitive to oxidative stress [52, 53]. Increased oxygenation and oxidative stress in the gut, often associated with dysbiosis (an imbalance in the gut microbiota), can negatively impact the abundance and survival of these sensitive anaerobic bacteria [52, 54]. This can further disrupt gut homeostasis. Our study clearly demonstrated that FLASH exposure which results less ROS production and oxidative stress maintains healthy gut microbiome compared to exposure with conventional dose rate. Our findings also highlighted the prevalence of some key microbiome species such as *Akkermansia muciniphila, Alistipes sp.* critical for mucosal epithelial regeneration and maintenance of barrier function [55, 56]. On the contrary Proton exposure with CONV dose rate promotes abundance of species such as *Romboitsia timonensis* which is associated with GI distress and inflammation [57]. In addition, our FMT study further validates that FLASH exposure favors the maintenance of functional healthy gut microbiome. FMT from FLASH exposed mice minimized the toxicity and improved survival in mice exposed to abdominal irradiation with Proton conventional dose rate.

## 4. Conclusion

This present study established a series of evidence suggesting the beneficial role of Proton FLASH abdominal exposure to spare mucosal toxicity at a dose level which is 100% lethal in conventional dose rate. Moreover, FLASH modulated the mucosal redox biology in favor of intestinal stem cell survival and stability of commensal microbiome to maintain gut homeostasis and regeneration. Importantly, this study reports for the first time the use of the IBA Proteus®ONE system to deliver pulsed, 228 MeV pencil beam scanning proton FLASH irradiation, thereby providing a unique platform for evaluating biological responses to this novel radiation modality. Therefore, this report supports that clinical application of Proton FLASH exposure will be safe and promise a higher therapeutic ratio for patients under abdominal radiotherapy.

## 5. Material and Method Animals

Seven to eight weeks-old male C57BL6/J mice, Lgr5-eGFP-IRESCreERT2 mice, Gt (ROSA)26Sortm4(ACTB-tdTomato-EGFP) Luo/J mice, and B6. Cg-Gt (ROSA)26Sortm9(CAGtdTomato)Hze/J mice were purchased (Jackson laboratories) for experimental studies. Lgr5-eGFP-IRES-CreERT2 mice were crossed with Gt (ROSA)26Sortm4(ACTB-tdTomato-EGFP) Luo/J mice (Jackson Laboratories) to generate Lgr5/eGFP-IRES-Cre-ERT2; R26-ACTB-tdTomato-EGFP [18, 58] and In Gt (ROSA)26Sortm4(ACTB-tdTomato-EGFP) Luo/J mice tdTomato is constitutively expressed (independent of Cre recombination) in the membrane of all cells, and therefore allows better visualization of cellular morphology. Lgr5-eGFP-IRES-CreERT2 mice were also crossed with B6. Cg-Gt (ROSA)26Sortm9(CAG-tdTomato) Hze/J mice (Jackson Laboratories) to generate the Lgr5-eGFP-IRES-CreERT2; Rosa26-CAG-tdTomato heterozygote for lineage tracing experiment. All the animals were maintained ad libitum, and all studies were performed under the guidelines and protocols of the Institutional Animal Care and Use Committee of the University of Kansas Medical Center. Animal studies were performed under the experimental protocols approved by the Institutional Animal Care and Use Committee of the University of Kansas Medical Center (ACUP number 22-01-215).

### Proton Irradiation procedure

Abdominal irradiation (ABI) was performed on mice that was anesthetized with Ketamine (87.5 mg/kg body weight) and Xylazine (12.5 mg/kg body weight) using a FLASH-capable proton synchrocyclotron (IBA Proteus^®^ONE) (Supplementary Figure 1). This is the first study to utilize this system to deliver pulsed, 228 MeV pencil beam scanning proton FLASH irradiation. A brass collimator was used to collimate the proton beams with a 2x3.5 cm^2^ opening which conferred dosimetric sparing to organs-at risk (OARs) outside of the mice’s gastrointestinal (GI) region. For this study, three proton ABI dose levels (14 Gy, 16 Gy and 18 Gy) were delivered at two FLASH (57.14-64.29 Gy/s and 43.21-49.38 Gy/s) and one CONV (0.5-1.0 Gy/s) dose rates to assess the differential impact of Proton FLASH and CONV irradiation survival outcomes. Post irradiation, all mice were monitored with activity levels and body weight assessed. Mice that reached a moribund status and/or experienced >20% body weight loss were considered endpoints and were euthanized.

The two different kinds of FLASH ABI fields were delivered: (1) High Uniformity/Low Ultra-High Dose Rate (L-UHDR) and (2) Low Uniformity/High Ultra-High Dose Rate (H-UHDR). H-UHDR was an earlier FLASH ABI iteration constructed from a 5x8 uniformly weighted proton spot map with 0.5 cm spot spacings centered on the brass collimator’s opening. However, proton-collimator scattering of the perimeter proton spots resulted in an increased central dose and relatively lower profile uniformities. Hence, L-UHDR was constructed from a 6x9 uniformly weighted proton spot map with similar spot spacings which extended into the brass collimator’s body. This extended spot map caused increased peripheral doses relative to central doses which improved 2D profile uniformities. However, L-UHDR dose rates were lowered relative to H-UHDR due to the increased number of spots delivered. Nominal beam delivery times were calculated with the nominal UHDR proton pulse structure of the Proteus^®^ONE synchrocyclotron, i.e. 1 kHz pulse repetition rate. All spot maps were designed with a raster-scanning pattern which eliminated spot-slewing deadtimes and this was verified retrospectively with log-file analyses. Subsequently, nominal FLASH dose rates were calculated by dividing the delivered FLASH ABI proton doses by their corresponding beam delivery times.

To ensure similar dose-rates across multiple FLASH ABI experiments, the number of pulses per spot was fixed. For 14, 16 and 18 Gy H-UHDR deliveries, 6, 7 and 7 proton pulses per spot were delivered respectively which corresponded to delivery times of 0.240 s, 0.280 s and 0.280 s respectively and correspondingly, FLASH dose-rates of 58.33, 57.14 and 64.29 Gy/s respectively. The differences in the dose-rates for the different dose levels are inevitable given the discreteness of the number of pulses delivered. For 14, 16 and 18 Gy L-UHDR deliveries, 6, 6 and 7 proton pulses per spot were delivered respectively which corresponded to delivery times of 0.324 s, 0.324 s and 0.378 s respectively and correspondingly, FLASH dose-rates of 43.21, 49.38 and 47.62 Gy/s respectively. Uniformity scores were defined as the ratio of the peripheral dose at 50% of the respective axes’ field sizes to the central field dose. HDR uniformities in the x- and y-directions were calculated to be 97.61% and 92.22% respectively whereas the corresponding ones for LDR were 98.26% and 95.50% respectively.

### Histology

The histological analyses jejunal sections were analyzed 72 hours post-irradiation with either LDR or HDR at 18 Gy and 21 Gy of Proton FLASH and CONV irradiation. Animals were euthanized and intestines were collected for histological analysis. The intestinal tissues were washed thoroughly by PBS to eliminate intestinal contents. Subsequently, the jejunum was fixed using 10% neutral-buffered formalin followed by embedding in paraffin and sectioning into 5-μm-thick sections for hematoxylin and eosin (H&E) and immunohistochemistry (IHC) staining. All the H&E staining was performed at the Histoserve Inc. (Germantown, MD, USA). The histopathological analysis was conducted for the assessment of crypt proliferation rate, villous length and crypt depth to analyze the intestinal tissue morphological changes.

### Crypt Proliferation Rate

To assess proliferative activity in villous epithelial cells, jejunal tissue samples were collected and subjected to paraffin embedding followed by immunohistochemical staining for Ki67. Paraffin-embedded sections underwent deparaffinization and rehydration through graded series of alcohols. Sections were then incubated overnight at room temperature with a monoclonal anti-Ki67 antibody (M7240 MIB-1; Dako). Nuclear staining was visualized using a streptavidin-peroxidase detection system in conjunction with diaminobenzidine (DAB) as the chromogen. A mild hematoxylin counterstain was applied to enhance tissue contrast and visualization. The identification of murine crypts within the tissue sections was employed using an established histological criterion as previously reported by Potten et al. [59]. To determine the proliferation index, Ki67-positive nuclei were expressed as a percentage of the total number of cells per crypt, with 50 well-defined crypts quantified per mouse.

### Determination of Villous Length and Crypt Depth

Quantitative analysis of villous height and crypt depth was performed in an unbiased, double-blinded manner using coded digital images obtained from H&E-stained intestinal sections. Measurements were conducted using ImageJ software (version 1.37). Crypt depth was assessed by measuring the distance, in pixels, from the crypt base to the crypt-villus junction, while villous height was measured from the crypt-villus junction to the tip of the villus. Pixel-based measurements were subsequently converted to micrometers using a calibration factor of 1.46 pixels per μm.

### Identification and Quantification of Lgr5⁺ Intestinal Stem Cells

To examine the survival of Lgr5+ crypt base columnar cells, Lgr5/eGFP-IRES-Cre-ERT2; R26-ACTB-tdTomato-EGFP mice were irradiated with ABI 21 Gy HDR of Proton FLASH and CONV irradiation. 96 hours of post-irradiation jejunal tissue were collected and processed for cryo-sectioning. To assess intestinal stem cell (ISC) survival, Lgr5⁺ crypt cells were identified based on enhanced green fluorescent protein (eGFP) expression, while overall crypt morphology and epithelial structure were visualized using tdTomato fluorescence expressed in all epithelial cell membranes. Quantification of Lgr5⁺ ISCs were performed by counting the number of eGFP⁺ crypts per field in a cross-section. For each mouse, a minimum of 10 randomly selected fields were analyzed.

### In-vivo Lineage Tracing Assay

To investigate the role of Lgr5-expressing ISCs in tissue regeneration under homeostatic conditions and in response to Proton FLASH and CONV radiation, an in-vivo lineage tracing assay was performed. Lgr5-eGFP-IRES-CreERT2 mice were crossed with B6. Cg-Gt (ROSA)26Sortm9(CAG-tdTomato) Hze/J mice (Jackson Laboratories) to generate the Lgr5-eGFP-IRES-CreERT2; Rosa26-CAG-tdTomato heterozygote mice. Mice were irradiated with ABI 16 and 18 Gy HDR of Proton FLASH and CONV irradiation, and tissue was harvested on day 6 post-irradiation. To initiate lineage tracing, adult mice were injected with tamoxifen (Sigma; 9 mg per 40 g body weight, intraperitoneally) in Cre reporter mice, allowing for the labeling of Lgr5+ lineages and their subsequent tdTomato (tdT)-positive progeny. This approach enabled the identification and tracking of Lgr5+ ISC lineages.

### Mouse Intestinal Crypt isolation and single-cell suspension

Mouse Intestinal tissue was isolated and washed three times with cold calcium- and magnesium-free Hank’s Balanced Salt Solution (HBSS) (Gibco, Cat no. 14170161). The cleaned tissue was open longitudinally and washed again 5 times with cold HBSS. After clearing all the intestinal content, tissue was cut into small fragments (approximately 3–5 mm in length) and incubated in 30 mM EDTA prepared in HBSS for 20 minutes on ice. Following incubation, the EDTA solution was discarded, and the tissue fragments were washed again in HBSS. To promote the release of epithelial cells vigorous shaking was performed (2–3 shakes per second for 3–5 minutes), followed by suspension was filtered through a 70 μm cell strainer. The resulting crypt-enriched fraction was centrifuged at 300 x g for 5 minutes at 4°C. Pelleted crypts were then enzymatically dissociated in 10 ml of Gentle dissociation buffer (StemCell technology) with gently shaking at rotatory shaker for 10 minutes at room temperature. Gentle pipetting was used to facilitate single-cell dissociation, and transfer the dissociated cells were mixed with 20 ml of ice-cold wash media containing DMEM/F12 media with 10µM of ROCK inhibitor and filtered using 40um cell strainer. The resulting cell suspension was centrifugation at 4 °C, 500 x g for 5 minutes and pelleted cells were resuspended in PBS with 2% BSA. Finally, cells were counted for viability check.

### Single-cell RNA sequencing

To characterize the epithelial cell populations at single-cell resolution to gain deeper insight into the regenerative and differentiation potential of the intestinal epithelium following 21 Gy of Proton FLASH and CONV irradiation, Single-cell RNA sequencing was performed. Mouse intestine was isolated 72 hours post-irradiation and single cell suspension was prepared as described above. The 10X Genomics Single Cell 3’ Expression v3.1 library preparation was performed using the 10X Genomics Chromium X. The single cells suspension was validated for viability and cell concentration using the Countess II FL Automated Cell Counter (Life Technologies) targeting ≥75% cell viability. The cell suspension was filtered through a FLOWMI Cell Strainer, 40µm (Thermofisher 50-136-7555) to yield a homogenous single cell suspension and adjusted to ∼1000 cells/µl to prepare cells for emulsification. The cell emulsification is performed with the 10X Chromium X using the Chromium Next GEM Single Cell 3’ GEM Library & Gel Bead Kit v3.1 (10X Genomics 1000120) and Chromium Next GEM Chip G Single Cell Kit (10X Genomics 1000127). The RT reaction to generate 10X barcoded single stranded cDNA, in the single cell containing GEMs, is performed using an Eppendorf MasterCycler Pro thermal cycler. The QC and quantification of the cDNA is determined using the Agilent TapeStation 4200 HS D5000 ScreenTape assay (Agilent 5067-5592). 3′ gene expression libraries were prepared using 10x Genomics protocols. Double-stranded cDNA was fragmented, end-repaired, A-tailed, and ligated with Illumina-compatible Unique Dual Index (UDI) adapters (Dual Index Kit TT Set A, 10X Genomics, 1000215), followed by indexing PCR. Libraries were validated on the Agilent TapeStation 4200 (D1000 ScreenTape, 5067-5582) and quantified using Roche LightCycler 96 with FastStart Essential Green Master Mix and KAPA Library Quantification Kit (KAPA, KK4903). Libraries were normalized to 2 nM, pooled, and sequenced on an Illumina NovaSeq X Plus using a 100-cycle kit (20104703) with a custom run format (Read1: 28 cycles, i7: 10 cycles, i5: 10 cycles, Read2: 90 cycles). Base calling, demultiplexing, and gene expression analysis were performed using Cell Ranger and Loupe Browser (10x Genomics).

### Single-cell RNA sequencing data processing and analysis

Single-cell RNA sequencing libraries were processed by aligning raw sequencing reads to the mm10 (mouse) reference genome and quantifying gene expression using Cell Ranger (v.7.1.0) with default parameters. Downstream analysis was performed on filtered features counts generated by Cell Ranger. Low-quality cells were removed using the following thresholds: number of genes per cell ≤ 250 or ≥ 9000, log10(Genes/UMI) < 0.8, or percent mitochondrial reads per cell > 10%. A gene was removed if it was not detected in at least 10 cells. Seurat (v.5.0.1) [60] was used to normalize samples with the log-normalize method and integrated using Harmony. The integrated dataset was then clustered using the Louvain algorithm and a resolution of 1. First, we identified epithelial and immune cell clusters using *Epcam* and *Ptprc*, respectively. The epithelial cells and immune cells were extracted to create two separate datasets. For each dataset set, we integrated and clustered the cells to further identify epithelial and immune cell populations. A resolution of 1 was used to cluster the epithelial cells and a resolution of 0.75 was used to cluster the immune cells. Epithelial cell clusters were annotated based on the expression of known markers as previously described [21]. The cell cycle phase was predicted for each cell using the CellCycleScoring function in Seurat. Differential gene expression (DGE) analysis was conducted for specific cell populations comparing the expression within a population between experimental groups (e.g., CBC cells: non-irradiated vs FLASH-irradiated). A Wilcoxon rank sum test was used to test for DGE, and a gene was considered significantly different if the adjusted p-value < 0.05, log2FC > 0.5 and proportion of cells expressing the gene > 0.1. Pathway over-representation analysis was conducted using the clusterProfiler package in R with the Reactome and KEGG pathways and MSigDb Hallmark gene sets. A pathway was considered significant if the p-value < 0.05. To predict the differentiation state of each cell, we used CytoTRACE [26], which outputs a stemness score ranging from 0 to 1. To have comparable stemness scores across all samples, we used the iCytoTRACE function in R.

### Metagenomic analysis

The mouse metagenomic analysis was performed on the mouse irradiated using ABI 18 Gy with both dose rates (LDR and HDR) of Proton FLASH and CONV irradiation. Fecal samples were collected at 24 and 72 hours in a sterile microcentrifuge tube and immediately snap freeze in liquid nitrogen. After that samples were kept at -80^0^C until sent for metagenomic analysis at CosmosID, Inc. (Germantown, MD, USA).

### DNA Extraction, Library Preparation and Sequencing

DNA from all the fecal samples were isolated using the QIAGEN DNeasy PowerSoil Pro Kit, according to the manufacturer’s protocol. Isolated DNA was quantified using Qubit Flex fluorometer and Qubit™ dsDNA Assay Kit (Thermofisher Scientific). DNA libraries were prepared using the Watchmaker DNA Library Prep Kit (7K0019-1K). Genomic DNA was fragmented using a master mix of Watchmaker Frag/AT Buffer and Frag/AT Enzyme Mix. IDT xGen UDI Primers and IDT Stubby Adapters were added to each sample followed by 7 cycles of PCR to construct the DNA libraries. The final DNA libraries were purified using CleanNGS magnetic beads (CleanNA) and eluted in nuclease-free water. Following elution, the libraries were quantified using the Qubit™ fluorometer dsDNA HS Assay Kit. Libraries were then sequenced on an Illumina NovaSeq X platform at 2x150bp. **Metagenomic Taxonomic Profiling Using Kepler**

Taxonomic profiling was performed using the patented Kepler multi-kingdom algorithm, which integrates three core components: (1) a pre-computed biomarker database (GenBook™), (2) k-mer–based taxonomic classification, and (3) probabilistic abundance estimation. In the database construction phase, high-quality microbial genomes across Archaea, Bacteria, Fungi, Protists, and Viruses were curated based on completeness, contamination, and assembly quality. Genomes were filtered to remove low-complexity regions, prophages, plasmids, and host contaminants. Genomic content was split into variable-length n-mers and structured phylogenetically, with shared and unique biomarkers mapped across a tree-like hierarchy. For classification, sample reads were decomposed into k-mers and matched against reference biomarkers from GenBook™, rapidly narrowing the candidate taxa list by excluding non-matching genomes. This enabled high-resolution taxonomic assignments down to sub-species level (ANI < 0.003). A refined abundance estimation was performed using a probabilistic Smith-Waterman algorithm on candidate taxa, allowing resolution of ambiguous reads through weighted assignment and iterative maximum likelihood estimation. This two-tiered approach ensured accurate, low-variance abundance estimates and high classification precision across microbial kingdoms.

### Functional Profiling of Metagenomic Reads

Metagenomic reads were quality-filtered and adapter-trimmed using BBduk [61]. High-quality reads were translated and aligned to the UniRef90 database (UniProt) [62], which clusters non-redundant protein sequences at ≥90% identity and ≥80% alignment coverage. Functional assignments were weighted by mapping quality, gene coverage, and sequence length to estimate community-wide gene family abundances, following the approach of Franzosa et al. (2018) [63]. Gene families were annotated to MetaCyc [64] reactions for reconstruction and quantification of metabolic pathways. Additional annotations included Enzyme Commission (EC) numbers, Pfam domains, CAZy enzymes, and Gene Ontology (GO) terms to provide a broad functional overview. Abundance values were normalized across samples using Total-Sum Scaling (TSS) to obtain "copies per million" (CPM), facilitating cross-sample comparisons.

### Quantitative Real time PCR

qPCR was performed to determine the WNT target gene and oxidative stress gene expression in intestinal epithelial cells following ABI of 21 Gy HDR using Proton FLASH and CONV irradiation. After isolation of epithelial cells as described above, total RNA was isolated from all the samples using RNeasy micro/mini kit from (Qiagen, Germantown, MD, USA). The concentration and purity of the extracted RNA were checked using a NanoDrop spectrometer (Thermo Scientific, Waltham, MA, USA). 1 µg of total RNA was reverse transcribed using RNA to cDNA EcoDry™ Premix (Double Primed) (Takara Bio USA Inc., San Jose, CA, USA). The reaction mixture was incubated for 1 h at 42°C; incubation was stopped at 70°C for 10 min. Quantitative real-time PCR (qPCR) was performed using the QuantStudio™ 7 Flex Real Time PCR System (Applied Biosystems™, New York, NY, USA) and SYBR Green Supermix (Bio-Rad, Hercules, CA, USA) with specific primers to the target genes in a 20 µL final reaction volume. GAPDH was used as a reference gene for sample normalization. The delta-delta threshold cycle (ΔΔCt) method was used to calculate the fold change expression in mRNA level in the samples. All the primer sequences used in this study are listed in Supplementary table 1. **Statistical analysis**

Mice survival/mortality was analyzed by Kaplan–Meier Log-rank (Mantel-Cox) test. For intestinal sampling regions were chosen at random for digital acquisition for quantitation. All quantitative data were analyzed using GraphPad Prism (v10.4.2). Comparisons between groups were made using unpaired two-tailed Student’s t-test. A p-value <0.05 was considered statistically significant. Data are presented as mean ± SEM. For scRNA-seq analysis, to test for differences in cell populations between Control, FLASH and CONV irradiation, we performed a Dirichlet-multinomial regression model since the cell proportions are compositional data. The models were fitted in R using the DirichletReg package with the alternative parameterization. To account for testing multiple comparisons, adjusted p-values using the Benjamini-Hochberg procedure. A cell population was considered significantly different between two conditions if the adjusted p-value < 0.05.

## Supporting information

Supporting Information contains supplementary figures and supplementary table.

## Authors contribution

S.S. conceived the project and designed the research; R.M.C., P.B. performed the experiments and interpreted the results; J.S., Y.L. performing the radiation, R.M.C., P.B., S.R., E.S., K.L.C. analyzed the data, R.M.C., S.S. wrote the first draft of the manuscript. R.M.C., P.B. E.S., K.L.C, S.S. edited the manuscript, S.K., D.K. R.C. providing the inputs.

S.S. provided essential reagents, tools and funding. All authors approved the manuscript.

## upplementary Figures

**Supplementary Figure 1:**
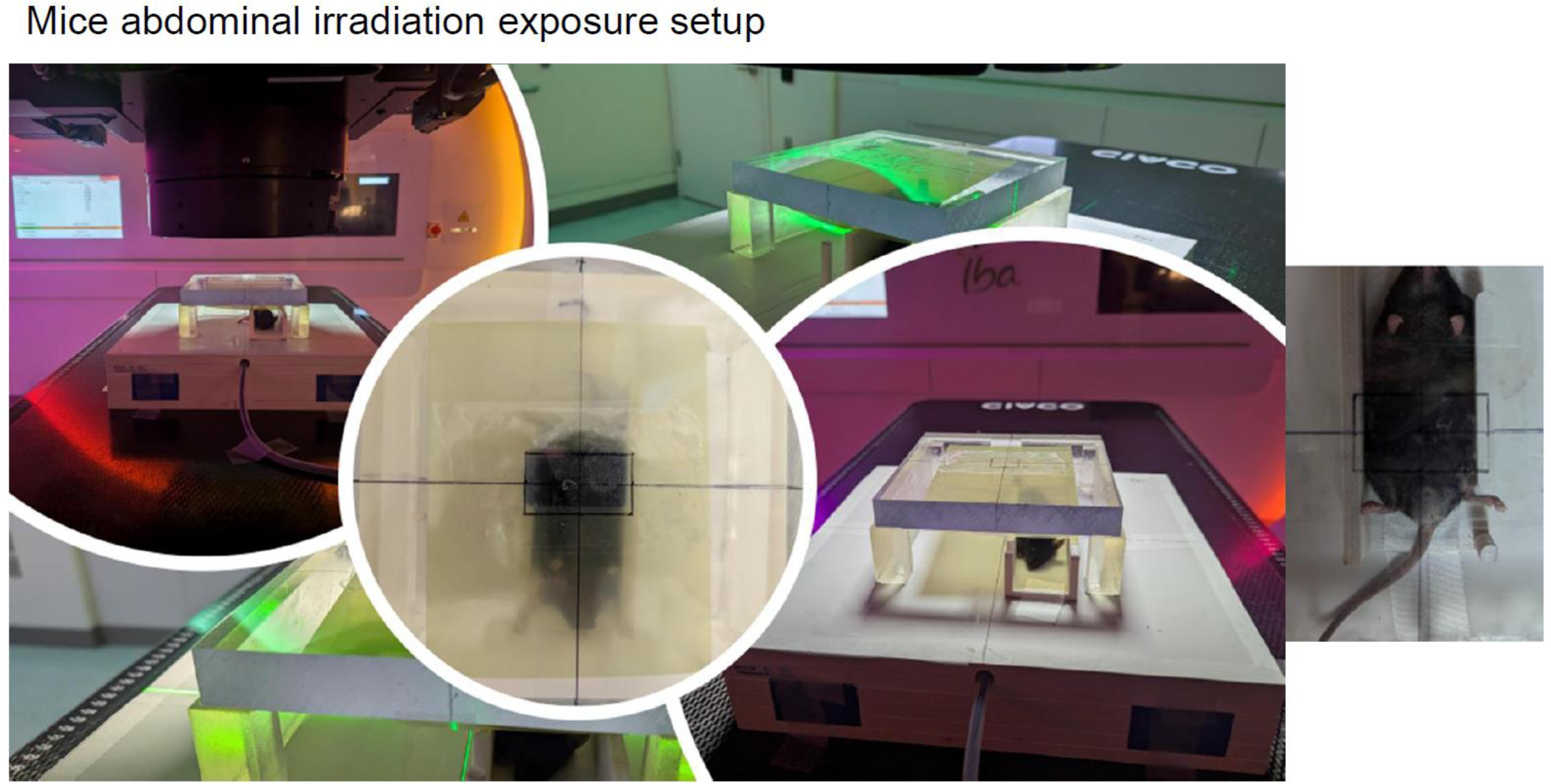
Mice abdominal irradiation exposure setup. Representative images showing the experimental setup for localized abdominal irradiation using Proton FLASH/CONV irradiation to mice. The mouse is placed in a custom-designed jig and only the abdominal region is exposed to radiation. Crosshair laser alignment is used for precise targeting of the abdominal field. Multiple views demonstrate the positioning of the mouse, and beam field alignment.

**Supplementary Figure 2:**
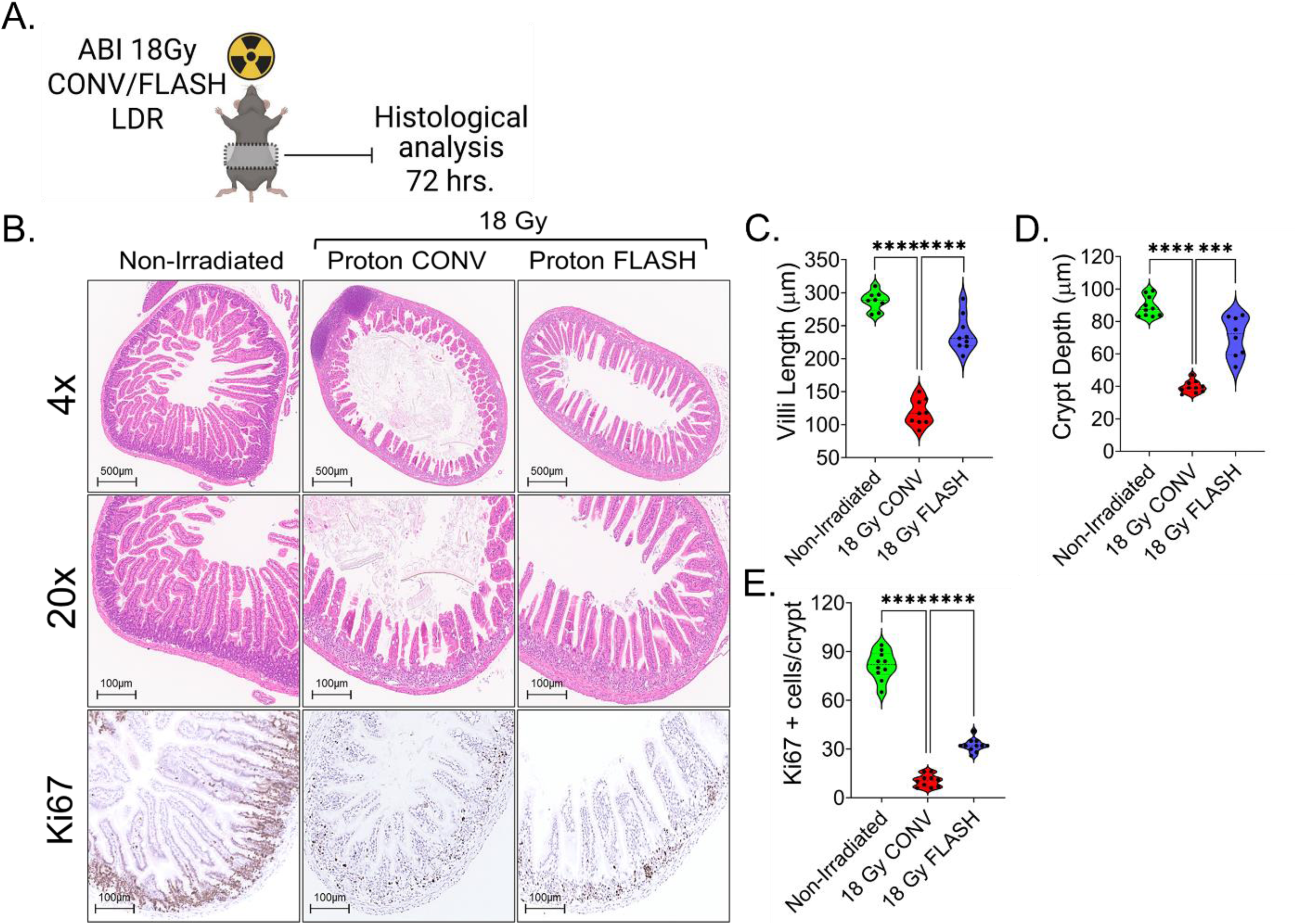
Proton FLASH irradiation preserves intestinal morphology following abdominal irradiation (ABI). (A) Schematic representation of experimental design. Mice received 18 Gy abdominal irradiation using either CONV or FLASH proton irradiation under LDR settings, followed by histological analysis of jejunal sections at 72 hours post-irradiation. (B) Representative H&E-stained jejunal cross-sections from CONV- and FLASH-irradiated mice under LDR conditions show improved mucosal preservation in FLASH-irradiated group (C, D) Violin plot demonstrating quantification of villus length and crypt depth. FLASH-irradiated tissues demonstrated significantly greater epithelial regeneration in compared to CONV IR. (B, E) showing the number of Ki67+ proliferative cells per crypt. FLASH-irradiated mice demonstrate significantly higher Ki67+ cell counts compared to CONV-irradiated animals. Data presented as mean ± SD, with Significant; ****p < 0.00005. n=10 mice per group.

**Supplementary Figure 3:**
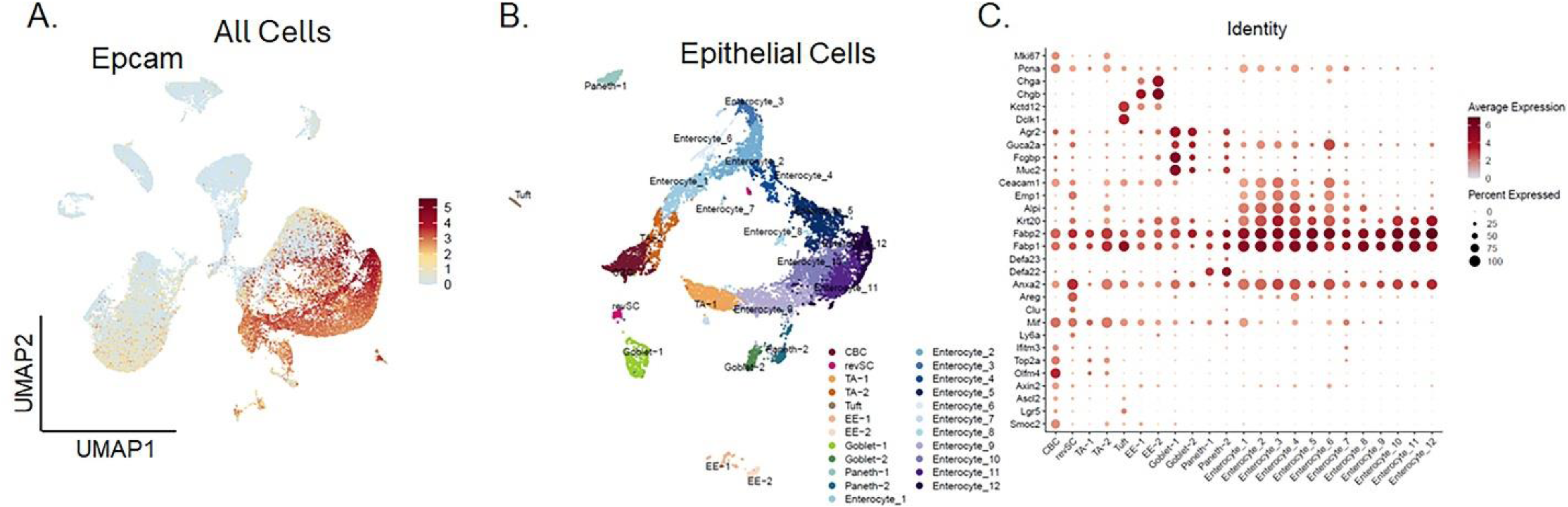
Single-cell transcriptomic analysis reveals different intestinal epithelial cell populations clusters. (A) UMAP shows Epithelial cells were identified by Epcam expression. (B)UMAP plot showing representation of 24 epithelial cell clusters identified from intestinal cells. (C) representation of Dot plot of selected marker genes across epithelial clusters, with dot size representing percentage of expressing cells and color indicating average expression.

**Supplementary Figure 4:**
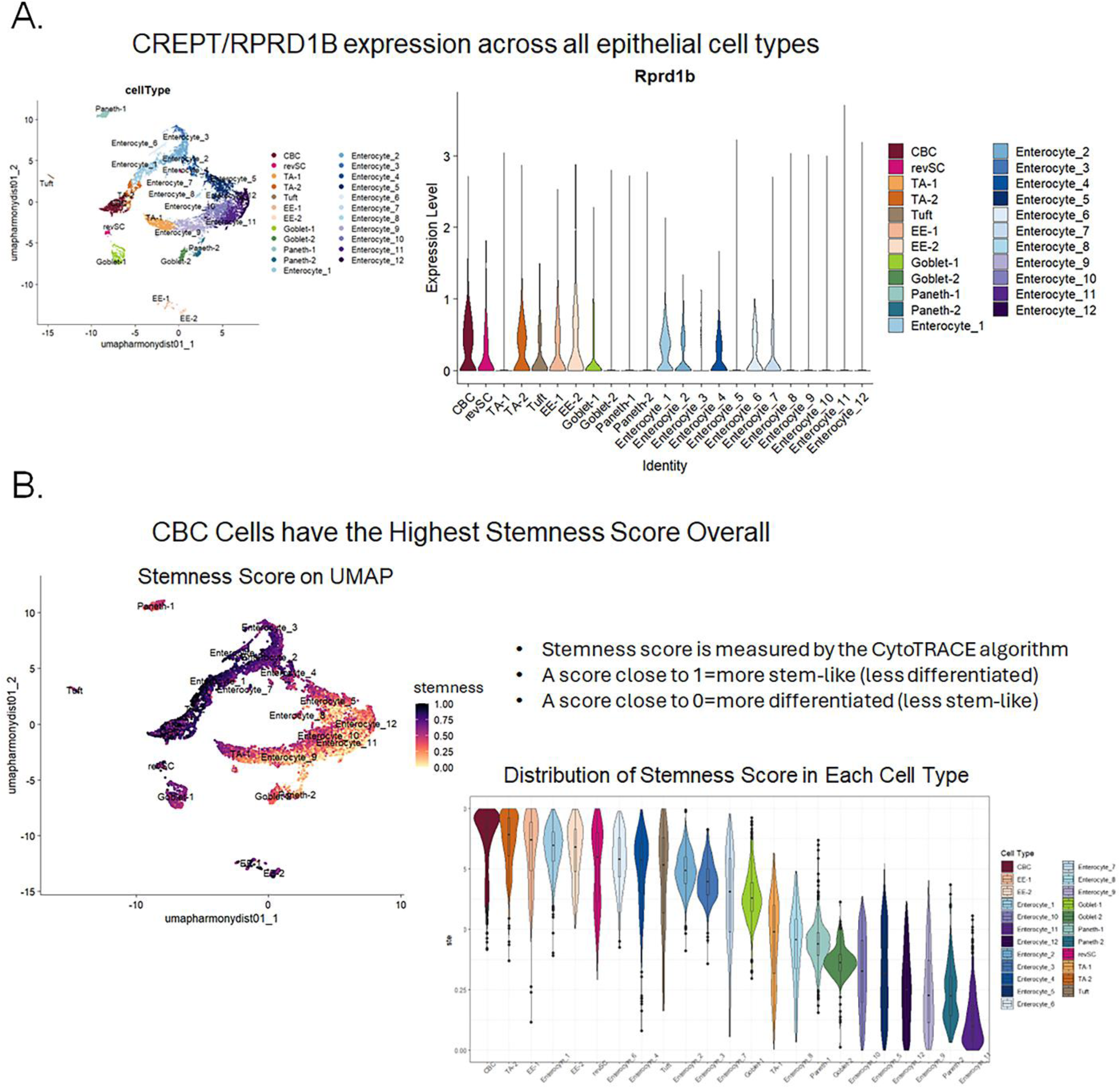
(A) Representation of CREPT/RPRD1B expression across all the epithelial cells clusters. (B) representing CBCs have highest Stemness score overall in all the epithelial cells clusters.

**Supplementary Table 1:**
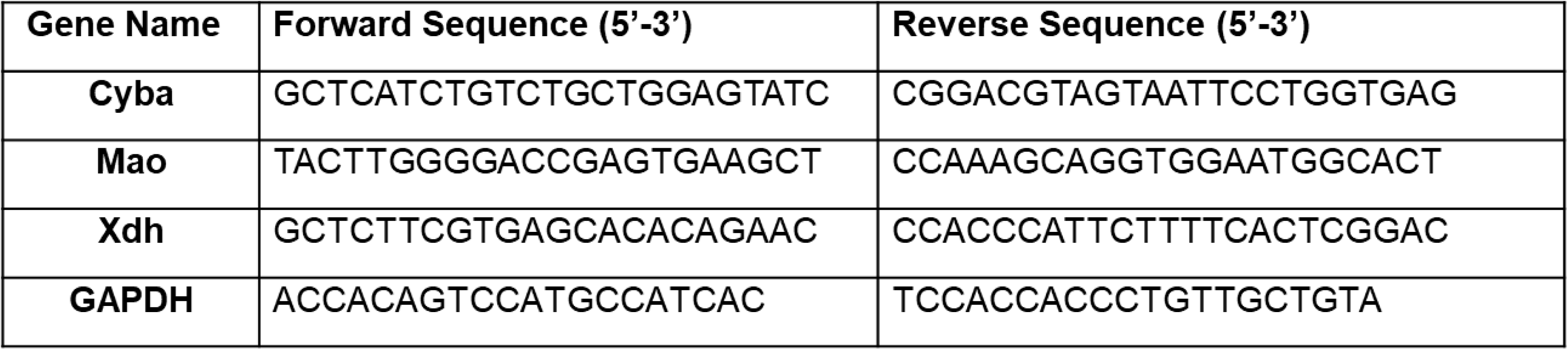
Primer Sequences.

